# Towards More Realistic Simulated Datasets for Benchmarking Deep Learning Models in Regulatory Genomics

**DOI:** 10.1101/2021.12.26.474224

**Authors:** Eva Prakash, Avanti Shrikumar, Anshul Kundaje

## Abstract

Deep neural networks and support vector machines have been shown to accurately predict genome-wide signals of regulatory activity from raw DNA sequences. These models are appealing in part because they can learn predictive DNA sequence features without prior assumptions. Several methods such as in-silico mutagenesis, GradCAM, DeepLIFT, Integrated Gradients and Gkm-Explain have been developed to reveal these learned features. However, the behavior of these methods on regulatory genomic data remains an area of active research. Although prior work has benchmarked these methods on simulated datasets with known ground-truth motifs, these simulations employed highly simplified regulatory logic that is not representative of the genome. In this work, we propose a novel pipeline for designing simulated data that comes closer to modeling the complexity of regulatory genomic DNA. We apply the pipeline to build simulated datasets based on publicly-available chromatin accessibility experiments and use these datasets to benchmark different interpretation methods based on their ability to identify ground-truth motifs. We find that a GradCAM-based method, which was reported to perform well on a more simplified dataset, does not do well on this dataset (particularly when using an architecture with shorter convolutional kernels in the first layer), and we theoretically show that this is expected based on the nature of regulatory genomic data. We also show that Integrated Gradients sometimes performs worse than gradient-times-input, likely owing to its linear interpolation path. We additionally explore the impact of user-defined settings on the interpretation methods, such as the choice of “reference”/”baseline”, and identify recommended settings for genomics. Our analysis suggests several promising directions for future research on these model interpretation methods. Code and links to data are available at https://github.com/kundajelab/interpret-benchmark.

## 1 Introduction

Complex machine learning models such as deep neural networks (DNNs) and support vector machines (SVMs) have demonstrated state-of-the-art performance at predicting genome-wide signals of regulatory activity as a function of the underlying DNA sequence [1, 2, 3, 4, 5]. When applying these models to decipher the complex *cis*-regulatory logic of the genome, it is important to understand which base pairs in an underlying genomic sequence are influential to the model’s prediction. In recent years, several interpretation methods such as in-silico mutagenesis (ISM) [6], GradCAM [7], DeepLIFT [8], Integrated Gradients [9] (for DNNs) and GkmExplain [10] (for SVMs) have been developed to answer this question. However, these methods have yet to be extensively benchmarked on models trained for regulatory genomic sequence prediction tasks, and our understanding of their relative drawbacks remains incomplete.

To evaluate the quality of the interpretations of models trained on regulatory sequence data, there are two main types of approaches. The first involves creating a completely simulated dataset by sampling motifs from position frequency matrices and implanting them uniformly into a randomly generated background sequence. This approach, which has been applied in several published works [8, 11, 12, 13] has the benefit of being in control of the exact ground truth. However, the problem with this approach is that these simulated sequences may lack the complexity of real biological data. Real protein binding is highly cooperative, with many factors often co-binding and affecting each other, and motifs also tend to exhibit positional preferences (for instance, motifs of activating transcription factors typically tend to occur near the center of a regulatory element). Conclusions drawn from these simplified datasets may not generalize to real data.

The second type of approach that has been used for benchmarking interpretation methods is to train models on real genomic sequences and use existing knowledge or heuristics to sanity-check the interpretation. For example, a user might look for relevant motifs that are known to exist in a cell type [2, 1, 5, 14], might verify that motif instances tend to occur within transcription factor footprints [5], might test for enrichment of disease-associated SNPs [15], and might even conduct empirical validation [5, 14]. While this approach uses real, representative data, it is difficult to gain insight into which patterns the interpretation method may have missed, as the ground truth is not definitively known.

### 1.1 Our Contributions

We devise a novel pipeline for generating simulated datasets to bridge the complexity gap between simple simulations and real genomic data. Using our pipeline, we generate 5 synthetic datasets based on chromatin accessibility data from 5 distinct cell types. On each of the 5 datasets, we train 2 different DNN models (pre-initialized using weights from DeepSEA Beluga [16] and Basset [2] respectively), as well as a gapped *k*-mer SVM [4]. We then benchmark 6 broad types interpretation methods (ISM, gradient-times-input, DeepLIFT, Integrated Gradients, GradCAM and GkmExplain) and explore multiple variations of DeepLIFT, Integrated Gradients and GradCAM, for a total of 19 different interpretation algorithms. Our findings can be summarized as follows:

1. We find that GradCAM-based methods, which were previously found to do well on a simplified simulated genomic dataset [11], perform poorly on our dataset (**Fig. 2**), particularly for the better-performing DeepSEA-like architecture that has shorter convolutional kernels in the first layer). In fact, we find that GradCAM performs worse than just using gradient-times-activation at the relevant layer (**Fig. 4**). We show this pitfall of GradCAM can be theoretically anticipated given the nature of genomic data (**Sec. 3.1**).
2. We investigate different settings for interpretation methods, such as the number of interpolation points for Integrated Gradients, the effect of different “references”/”baselines”, and the benefit of using the ‘RevealCancel’ variant of DeepLIFT (**Sec. 3**). We identify recommended settings for practitioners.
3. We occasionally find that Integrated Gradients (IG) performs worse than gradient-times-input (**Fig. 2**). Consistent with the findings of Jha et al. [17], we speculate that this is due to IG’s linear interpolation path, as interpolated inputs can be out-of-distribution. We show empirically that linear interpolation between the reference and target output can cause the output logits to behave erratically (**Fig. S10**).
4. We find that for the poorer-performing Basset-initialized model, GkmExplain importance scores perform close to the best deep-learning-based importance scores, showing the importance of training a SVM as a strong baseline alongside deep learning models (**Fig. 2**).

## 2 Methods

### 2.1 Description of Interpretation Algorithms Used

We provide a brief overview of the interpretation methods considered in this work. For all these methods, the “output” we used was the logit of the sigmoid (i.e. the log-odds probability of the positive class).

**In-Silico Mutagenesis (ISM)** quantifies the importance of individual bases by making *in-silico* perturbations to individual bases in the input and observing the change in the output. In our implementation, we first computed the delta in the output logit when the base at a given position is mutated to the 3 other possible bases, and then defined the importance of the position as the negative of the average delta (thus, if mutating a position tends to decrease the logit, the position will be assigned a positive importance). ISM is attractive as it is a faithful depiction of how the model responds to a perturbation. A key drawback is that it only reflects the impact of making a single perturbation; if the output has saturated in terms of its sensitivity to an input feature, then perturbing that feature may not affect the output. Another drawback is that the output has to be recomputed every time there is a perturbation, though faster implementations exist [18].

**Gradient-times-input** [19] does an elementwise multiplication of each input feature with the gradients of the output on the input features. In the context of genomics, where inputs are one-hot encoded, this amounts to computing the gradient on the bases that are present in the sequence. The advantage of this approach is its simplicity; the disadvantage is that gradients can saturate [8], and can thus miss important features.

**DeepLIFT** [8] is an explanation method for DNNs devised to counter the saturation faced by gradient-times-input, while retaining some computational efficiency. DeepLIFT works by comparing the activations of neurons on the actual input to the activations of the neurons on a “reference” or “baseline” input, and backpropagating an importance signal (“contribution scores”) in such a way that the sum of contributions across all input features will equal the difference of the output activation from its reference value. A key drawback of DeepLIFT is that it is a heuristic importance-scoring method backpropagates the importance signal based on the network’s internal wiring; thus, two identical models with different internal wiring could, in principle, produce different outputs (i.e. DeepLIFT is not “implementation invariant”).

A note on different variants of DeepLIFT that exist in the literature: the original DeepLIFT paper proposed two rules for backpropagating importance through a nonlinear activations: the **“Rescale” rule** and the **“RevealCancel” rule**, where the latter rule was designed to reveal cases where different inputs to a neuron may have canceled each other out (thereby superficially making it appear as though the neuron did not receive any significant inputs). The authors recommended that, for genomics applications, the RevealCancel rule be applied to fully-connected layers and the Rescale rule be applied to convolutional layers. Since the publication of the DeepLIFT paper, several additional repositories have implemented DeepLIFT in a more extensible way than was done in the original repository [20, 21, 22] - however, at the time of writing, these implementations do not support the RevealCancel rule and instead use Rescale at all activation layers. Prior work has not investigated how much benefit is conferred by the RevealCancel rule on real genomic data, particularly when shuffled sequences (discussed below) are used as a reference.

Like DeepLIFT, the method of **Integrated Gradients** (IG) [9] also uses a reference-based approach to address the saturation problem faced by gradient-times-input. IG averages the input feature gradients over several pseudo input examples that are generated by linearly interpolating between the reference example and the actual input example, and then multiplies these average gradeints elementwise with the difference of the actual input features from their reference values. Because IG only relies on model gradients, it is guaranteed to produce identical outputs for functionally equivalent models irrespective of the internal network implementation - i.e. it is “implementation invariant”. However, although it is implementation invariant, it is not invariant to the choice of interpolation path; this is discussed more in **Sec. 3.2**. Another pitfall of integrated gradients is computational efficiency, as the runtime is proportional to the number of interpolation points (though it should be noted that this runtime can be reduced with sampling approaches [23]).

The **choice of “reference”** is of key importance when using a method like DeepLIFT or IG [24]. In the original DeepLIFT paper [8], the authors used a reference representing the expected base frequency; for example, with 40% GC content and base ordering of ACGT, this reference would look like a *L*× 4 vector where *L* is the sequence length and each column is [0.3, 0.2, 0.2, 0.3]. Another commonly-used reference is the all-zeros reference, which is the default reference in most implementations of DeepLIFT/IG and was the only reference explored in the original IG paper [9]. However, the supplement of the DeepLIFT paper showed strong performance using a collection of reference sequences derived by dinucleotide shuffling the original sequence; in this case, contribution scores are averaged across each reference in the collection. Prior work has not benchmarked the use of shuffled references extensively. It should be noted that using multiple references adds computational burden proportional to the number of references.

**GradCAM** [7] is an explanation method that was developed for computer vision data. When applied to the sequential inputs used in genomics, the method works as follows: for a given target output task *y*^*o*^, and a given convolutional layer in the network (the GradCAM paper recommends using the last convolutional layer), a set of weights 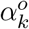 are determined for each channel *k* in the convolutional layer (the GradCAM paper refers to channels as “feature maps”) by averaging gradient of the output over all positions in the channel; thus, if 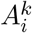 represents the activation of channel *k* at position *i*, then 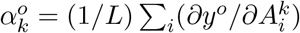, where *L* is the total length of the conv layer. An importance vector is then computed as 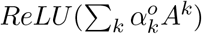; this vector has length *L*, and is projected onto the input sequence via linear interpolation (the original GradCAM paper uses bilinear interpolation as it deals with image-like inputs).

**Zheng et al. [11]** used a variant of GradCAM where the importance vector was computed at the first conv layer rather than the last. According to the [code](https://github.com/Pandaman-Ryan/AgentBind/blob/f5ebe339960b9627328091e9825fb5ffaff47110/AgentBind-Release-Version-DanQ/model/visualize_weights.py#L45) released by the authors, this importance vector was mapped onto the input sequence using a different method than the linear interpolation of GradCAM: the importance of a sequence position was defined to be the average importance over all conv neurons whose receptive field overlapped the position. Finally, the sequence-level importance was multiplied elementwise with the input gradients for finer base-pair-level resolution. Because it is a nonstandard version of GradCAM, we refer to this approach as the “Zheng et al.” method.

**GkmExplain** [10] is an explanation method for support vector machines (SVMs) trained on genomic sequence data with gapped *k*-mer kernels. GkmExplain decomposes the output of the SVM into contributions from individual bases, and was shown to produce state-of-the-art results at identifying motifs in regulatory DNA sequences. It is orders of magnitude more computationally efficient than applying ISM to SVMs.

### 2.2 Simulation Pipeline

Here is the workflow we used for generating a simulated datasets (summarized in **Fig. 1**):

**Figure 1:**
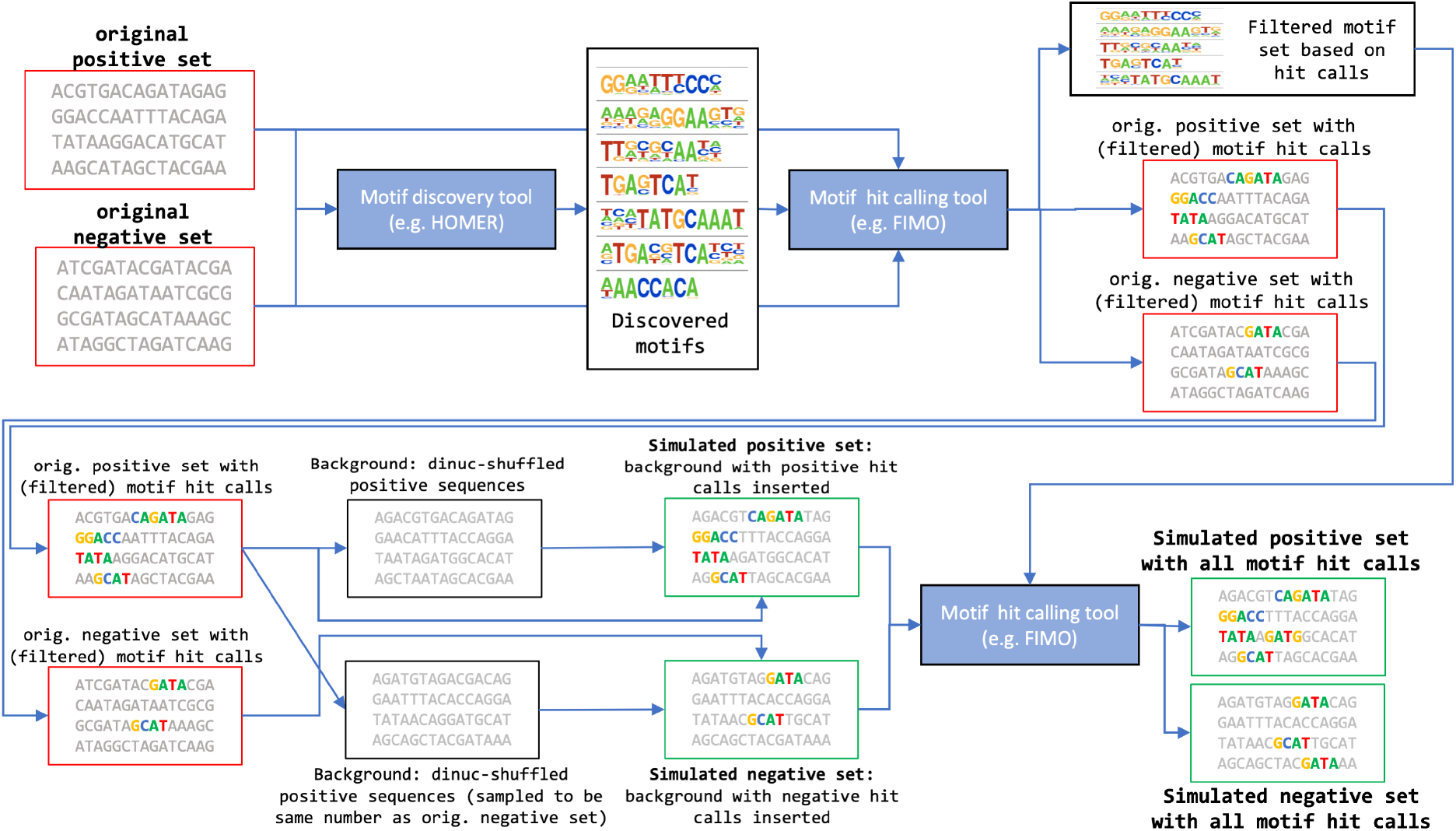
Summary of our simulation pipeline.

1. Given a set of positive and negative sequences (e.g. identified through functional genomic assays), run a motif discovery algorithm (in our case, HOMER) on the positive set, using the negative set as a background. This will identify the enriched motifs in the positive sequences.
2. Run a motif calling algorithm (in our case, FIMO) on *both* the positive and negative set using the motifs discovered in the previous step, in order to find instances of these motifs in both the positive and negative sets. It is important to keep track of motif instances in the negative set as negative sets in real genomic data do contain instances of motifs that may be enriched in the positive set (they are often distinguished from motif instances in the positive set by positional or motif co-binding patterns). Based on the motif calling statistics, filter out any motifs that do not pass a particular enrichment threshold. Details about the enrichment thresholds we used are in **Sec. S1.1**.
3. For each positive sequence, dinucleotide shuffle the sequence to randomize it, and implant the motif instances (identified in the previous step) at their known original locations in the sequence (note that the original motif match is implanted; we do not sample from the motif PWM). The dinucleotide shuffling is intended to scramble undetected motif instances hidden at unknown positions in the sequence.
4. GC imbalance between positive and negative sets can sometimes confound interpretation; to eliminate this imbalance, for each negative sequence we randomly choose a sequence from the positive set and dinucleotide-shuffle it in order to form the “background” for the negative sequence. Then, as is done for the positive set, we insert motif matches identified by the motif calling algorithm in the negative set at their original locations in the negative sequence.
5. Run the motif calling algorithm again on the positive and negative sequences in order to identify both the implanted motif instances as well as motif instances that may have spontaneously appeared in the dinucleotide-shuffled background sequences.

The advantages of this approach are: **(1)** any motifs that were not discovered by motif discovery are scrambled by dinucleotide shuffling; thus, we have some notion of ‘ground truth’ features; **(2)** because the background is similar for the positive and negative sets, we should not expect interpretation to be confounded by GC bias; **(3)** because motifs are retained at their original positions, and motif instances can be present in the negative set too, we challenge the model to learn complex, co-operative TF binding patterns to discern the positive set.

### 2.3 Datasets

In this work, we generated our simulated positive set based on 400 bp centered around the summits of ENCODE [25] IDR [26] peaks accessible in one of the Tier 1 cell lines A549, GM12878, H1ESC, HepG2, or K562 (accession numbers ENCSR149XIL, ENCSR000ELW, ENCSR794OFW, ENCSR000EOT, and ENCSR000EMT). For our negative set, we compiled a union of IDR peak regions that were accessible in any cell type and used 400bp regions centered around the summits of these peaks, filtering out regions that overlapped with peaks from the cell type used as the positive set. Sequences with nonstandard bases (i.e. non ACGT) were also filtered out. We delegated 20% of the simulated sequences as a testing set.

To sanity check that our datasets contains non-random motif co-occurrence patterns, we visualized a heatmap of motif-cooccurence for the A549 cell type in **Fig. S7**. To verify that our dataset contains motif positional preferences, we visualized the positional distribution for two A549 motifs in both the positive and negative set in **Fig. S8**; these distributions show that both motifs are more likely to occupy the center of the 400bp regions in the positive set as compared to the negative set.

### 2.4 Model Training

For each cell type, we pre-initialized two models using publicly available weights from DeepSEA Beluga and Basset respectively, and fine-tuned the models on our simulated data. For both models, the DeepSEA Beluga and Basset architectures were adapted to be single-task architectures (we initialized the weights for the output node in the single-task model using the weights for the corresponding output task in the pre-trained multitasked models), and the available weights were also trimmed to accomodate our 400 bp input.

During training, we used a custom data loader that sampled each batch to contain 50% positives and 50% negatives. We used the hyperparameters of learning rate = 0.001, batch size = 300, loss = binary cross-entropy, optimizer = Adam, and maximum number of training batches = 30,000. Early stopping was done by evaluating the auROC on a fixed sample of 2,500 positive and 2,500 negative sequences from the test set every 100 batches, and pausing if there was no improvement over 3 consecutive evaluations (300 batches). The auROCs and auPRCS achieved by the model on the full test set for each cell line are listed in **Tab. 1**. The numbers of positives and negatives sequences are listed in **Fig. S6**.

**Table 1:**
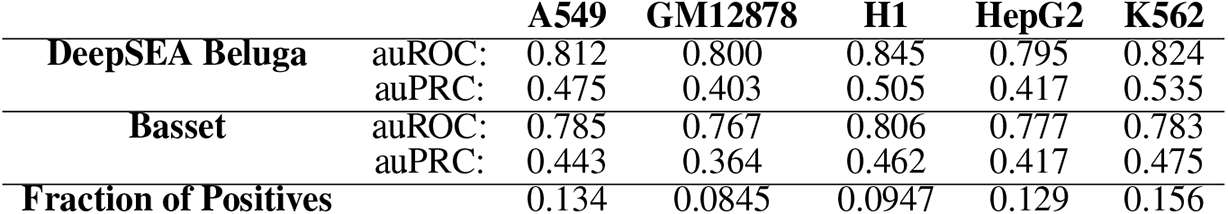
Test auROC/auPRC for each finetuned model and cell line.

For our baseline gkm-SVM models, we used the default settings from the lsgkm package [4] (wgkm kernel with input length *L* = 11, k-mer size *k* = 7, and gaps *d* = 3).

### 2.5 Evaluating Interpretation Methods

To calculate an auPRC for motif recovery, we follow Koo and Ploenzke [13] and define positive positions as sequence positions that fall within motif regions and negative positions as sequence positions that fall within nonmotif regions. For a given cell line and interpretation method, the auPRC for distinguishing positive positions from negative positions in a single sequence is found by ranking the positions in descending order of the absolute value of importance. Due to the computational overhead involved with running some interpretation methods, we limited our evaluation to the top 10K training-set sequences in each cell line that were most strongly predicted as positive by the finetuned DeepSEA Beluga model. In our barplots, we show the mean auPRC across all sequences and report the error bars as the standard error of the mean auPRC.

## 3 Results

An overall comparison of different types of interpretation methods is shown in **Fig. 2**, and importance scores at example sequences are visualized in **Fig. 5, S14 & S13**. A few points of interest:

**Figure 2:**
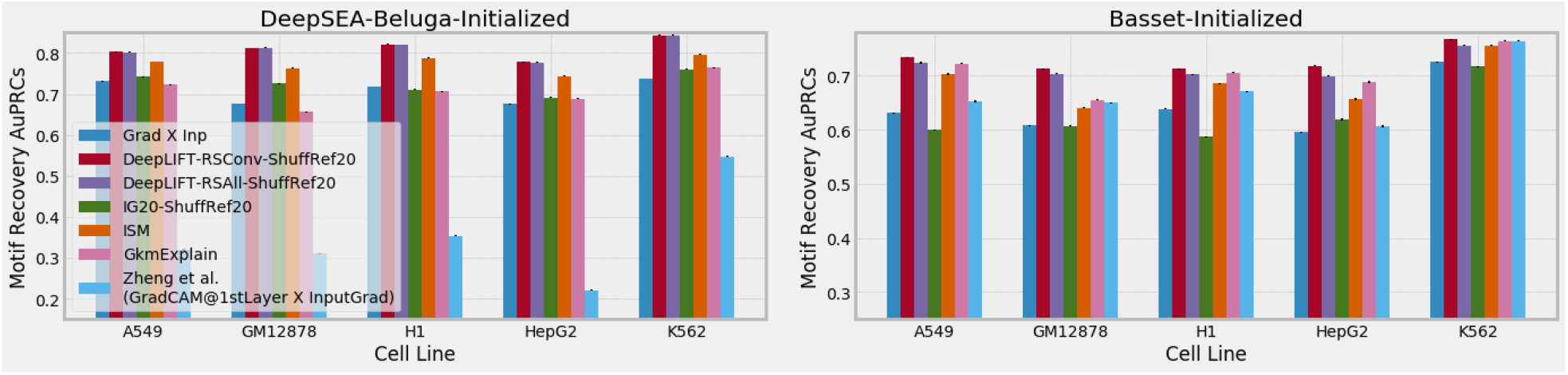
Overall omparison of different interpretation methods. Left panel shows scores from deep learning models initialized with DeepSEA-Beluga weights and fine-tuned on the respective simulated data; right panel used Basset-initialized weights. GkmExplain scores in both panels are identical as they are derived from the SVMs. ‘IG20’ denotes Integrated Gradients with 20 interpolation points. ‘ShuffRef20’ indicates 20 dinuc shuffled references were used per sequence. For DeepLIFT, ‘RSAll’ indicates the variant with the Rescale rule applied at all layers, and ‘RSConv’ indicates the variant where Rescale is only applied to the conv layers. See **Sec. 2.1** for more information on the methods.

1. We find that DeepLIFT and ISM tend to perform best and second-best; surprisingly, ISM performed below DeepLIFT, which we discuss more in **Sec. 4**.
2. The GradCAM-based method of Zheng et al. [11] performed relatively poorly in this simulation, particularly for the DeepSEA-like architecture (which has shorter kernels in the first convolutional layer); we explore this in **Sec. 3.1** and **Fig. 4**.
3. IG sometimes performed worse than grad-times-input; we discuss this in **Sec. 3.2**.
4. We verified that the poorer performance of IG was not due to an insufficient number of interpolation points; **Fig. S9** shows that Integrated Gradients with 20 interpolation steps and 10 interpolation steps gives nearly identical performance.
5. We find that DeepLIFT with multiple dinuc-shuffled references consistently performs better than an all-zeros base frequency reference (**Fig. 3**). We also saw that DeepLIFT performance improves with more shuffled references, tapering after 20.
6. The Rescale-only variant of DeepLIFT performed similarly to the variant with Rescale on the conv layers and RevealCancel at the dense layers (with the latter variant doing slightly better on Basset-initialized models). We note that this difference is small compared to the choice of reference (**Fig. 3**).
7. GkmExplain interpretations performed well compared to interpretations from the poorer-performing Basset-initialized model (**Tab. 1**), demonstrating the value of training SVMs as a baseline.

**Figure 3:**
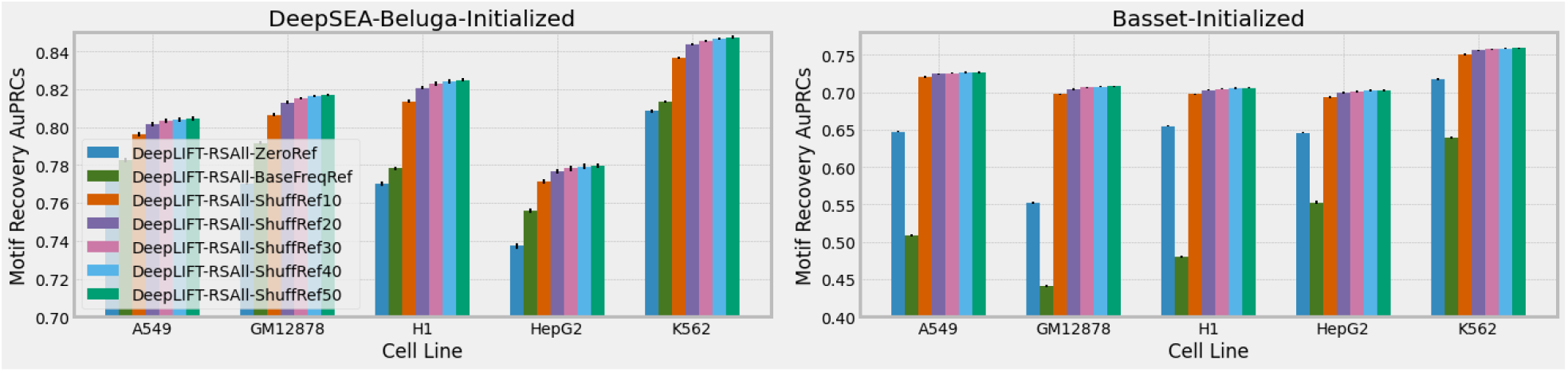
Comparison of different DeepLIFT references. ‘ZeroRef’ is an all-zeros reference, BaseFreqRef is a reference with fractional values representing expected base frequences, ShuffRefX indicates dinuc shuffled references, with X denoting the number of references used per sequence. Refer to **Sec. 2.1** for more information.

### 3.1 GradCAM-like methods perform relatively poorly, but attributions at convolutional layers show promise

When investigating GradCAM-like methods, we began by benchmarking the version of GradCAM commonly used in computer vision: importance was computed at the last conv layer and mapped to the input using linear interpolation. We found that this achieved poor performance (**Fig. 4**). To investigate whether this was because the spatial resolution of the last conv layer was too coarse for genomics, we also computed GradCAM scores using the 1st conv layer, and further explored the method of Zheng et al. [11] (which multiplied GradCAM-derived scores from the 1st layer with the input gradients); in both cases, we used the mapping method of Zheng et al. [11] to map the conv layer scores onto the input, as we found it performed better at the 1st layer. However, these approaches still achieved poor performance in our simulation (**Fig. 4**), particularly for the DeepSEA-like architecture that has shorter convolutions in the first layer. We argue these findings are to be expected based on the weighting approach used by GradCAM:

**Figure 4:**
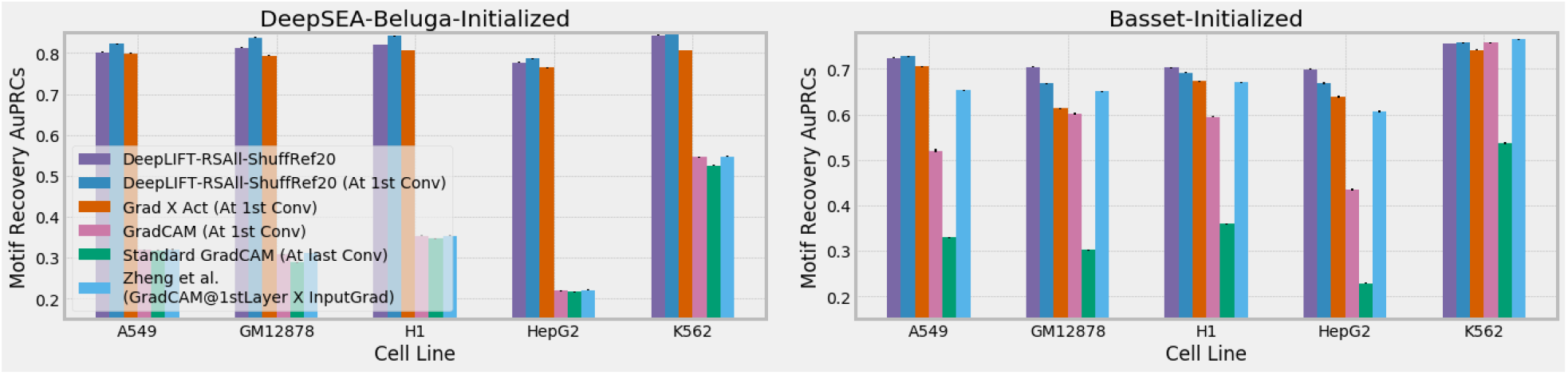
Comparison of GradCAM and related methods. ‘DeepLIFT-RSAll-Shuff20’ is the same bar as displayed in **Fig. 2**. See **Sec. 2.1** **&** **3.1** for more info on the methods.

Recall from **Sec. 2.1** that GradCAM computes “weights” 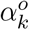 for each convolutional channel *k* by averaging the gradient of the output *y*^*o*^ over all positions in the channel. By using the same weight for all neurons in a channel, GradCAM assumes that an activated neuron in the layer has equal importance to the output task regardless of both the *position* at which the neuron is activating, as well as the *context* of which other neurons in the layer are activating nearby (this loss of information is illustrated in **Fig. S11 & S12**). In genomics, where motifs can have different effects depending on whether they occur at the center vs the flanks, and also co-bind with other motifs, the assumption of positional and contextual independence may not be valid. In fact, experiments in computer vision have reported that this aspect of GradCAM leads it to fail “anywhere besides the last convolutional layer” [27], and the GradCAM paper itself states that “localization becomes progressively worse as we move to shallower convolutional layers. This is because the later convolutional layers capture high-level semantic information and at the same time retain spatial information, while the shallower layers have smaller receptive fields and only concentrate on local features that are important for the next layers” (section 5.2 of the [supplement](https://openaccess.thecvf.com/content_iccv_2017/html/Selvaraju_Grad-CAM_Visual_Explanations_ICCV_2017_paper.html)). This perspective is consistent with the fact that GradCAM performs worse on the DeepSEA-like architecture compared to the Basset-like architecture; the DeepSEA-like architecture has convolutions of length 8 in the first layer, while the Basset-like architecture has convolutions of length 19 in the first layer - thus, individual neurons in the first layer of a Basset-like architecture are more capable of recognizing complete motifs on their own. In other words, when longer filters are used in the first convolutional layer, the importance of a neuron in the first layer is less likely to depend on the context of which other neurons in the layer are firing nearby.

While it may be tempting to conclude from this that it is safe to use GradCAM if we confine ourselves to architectures that have longer filters in the first layer, doing so would come at the expense of model performance; trends in both computer vision [28] and in regulatory genomics [29] have found that performance can be improved by “factorizing” layers that have large convolutional kernels into several layers that have smaller convolutional kernels. While it is tempting to think that we can safely trade of model performance in exchange for improved interpretability, the reality is that the two are not completely independent; for example, in our study, we found that the best explanations across all methods were derived from the better-performing DeepSEA-initialized models (**Fig. 2, Tab. 1**).

A question that may arise is why the benchmarking of Zheng et al. [11], which used a DeepSEA-like architecture, did not identify the GradCAM-based method as a poor-performing method. We suspect this is due to the fact that Zheng et al. [11] benchmarked their method on a variation of the simplified simulation used in Shrikumar et al. [8], where the positive set was identifiable by the presence of a TAL1 motif that was randomly inserted into a 1kb sequence. Because the core TAL1 motif is 6bp long (CAGATG), and the model was not required to distinguish TAL1 from other very similar motifs, it is possible that individual convolutional filters in the first layer of the DeepSEA-like architecture were fully capable of recognizing the TAL1 motif; thus, the model in this simulation was not required to learn complex patterns involving muliple neurons activating together, allowing GradCAM to perform well. This example highlights the importance of using realistic simulations to benchmark methods.

As an alternative to GradCAM-based methods, we explored what would happen if we did not sacrifice any positional information, but rather simply computed standard neuron attributions at the same layer as GradCAM. We first calculated both gradient-times-activation (grad-times-act) and DeepLIFT scores directly at the first conv layer, and then transformed these conv-layer scores to the dimensions of the input by using an analogous method to the one used in Zheng et al. (i.e. we first summed the conv-layer scores along the channel dimension to get a vector of length equal to the length of the conv layer, and then for each input position we averaged the importance over all conv-layer positions whose receptive field overlapped the input position; we did not discard negative scores). Grad-times-act-at-1st-conv substantially outperformed GradCAM-at-1st-conv-layer by very large margins for the DeepSEA-like architecture. For the Basset-like architecture, grad-times-act-at-1st-conv outperformed GradCAM-at-1st-conv-layer by large margins on 3 cell types (A549, H1 and HepG2), outperformed it by a small margin on 1 cell type (GM12878), and underperformed it by a small margin on 1 cell type (K561). DeepLIFT-at-1st-conv-layer consistently outperformed grad-times-act-at-1st-conv-layer, which is to be expected given that DeepLIFT was specifically designed to overcome the limitations of using just gradients.

One striking finding was that DeepLIFT at the 1st conv layer sometimes outperformed DeepLIFT at the input layer (blue vs. purple bars in **Fig. 4**), suggesting there is promise to computing importance scores at layers other than the input layer. We discuss this more in **Sec. 4**.

### 3.2 On the performance of Integrated Gradients

We found that IG sometimes performs worse than grad-times-input (**Fig. 2, 5 & S14**), even though IG was designed to combat the saturation problem faced by grad-times-input. In **Sec. 2.1**, we noted that although Integrated Gradients is “implementation invariant”, it is not invariant to interpolation path that is used between the reference and the input. By default, the linear interpolation path is used - however, such a path can produce out-of-distribution examples during interpolation. In the case of genomic data, interpolating between two one-hot encoded inputs results in an input containing fractional values, which the model has never seen during training. Jha et al. [17] demonstrated that the linear interpolation of IG performed poorly in the context of splicing codes, though to our knowledge prior work has not demonstrated that it is a drawback for regulatory genomic data as well. To support the hypothesis that linear interpolation can cause issues, we plotted histograms of the sigmoid logits during interpolation between original and reference A549 sequences in **Fig. S10**. We found that the logits on interpolated inputs do not smoothly transition from the reference logits to the actual logits; indeed, they can move further away from the actual logits during interpolation, confirming that non-one-hot-encoded inputs can produce unexpected model outputs.

**Figure 5:**
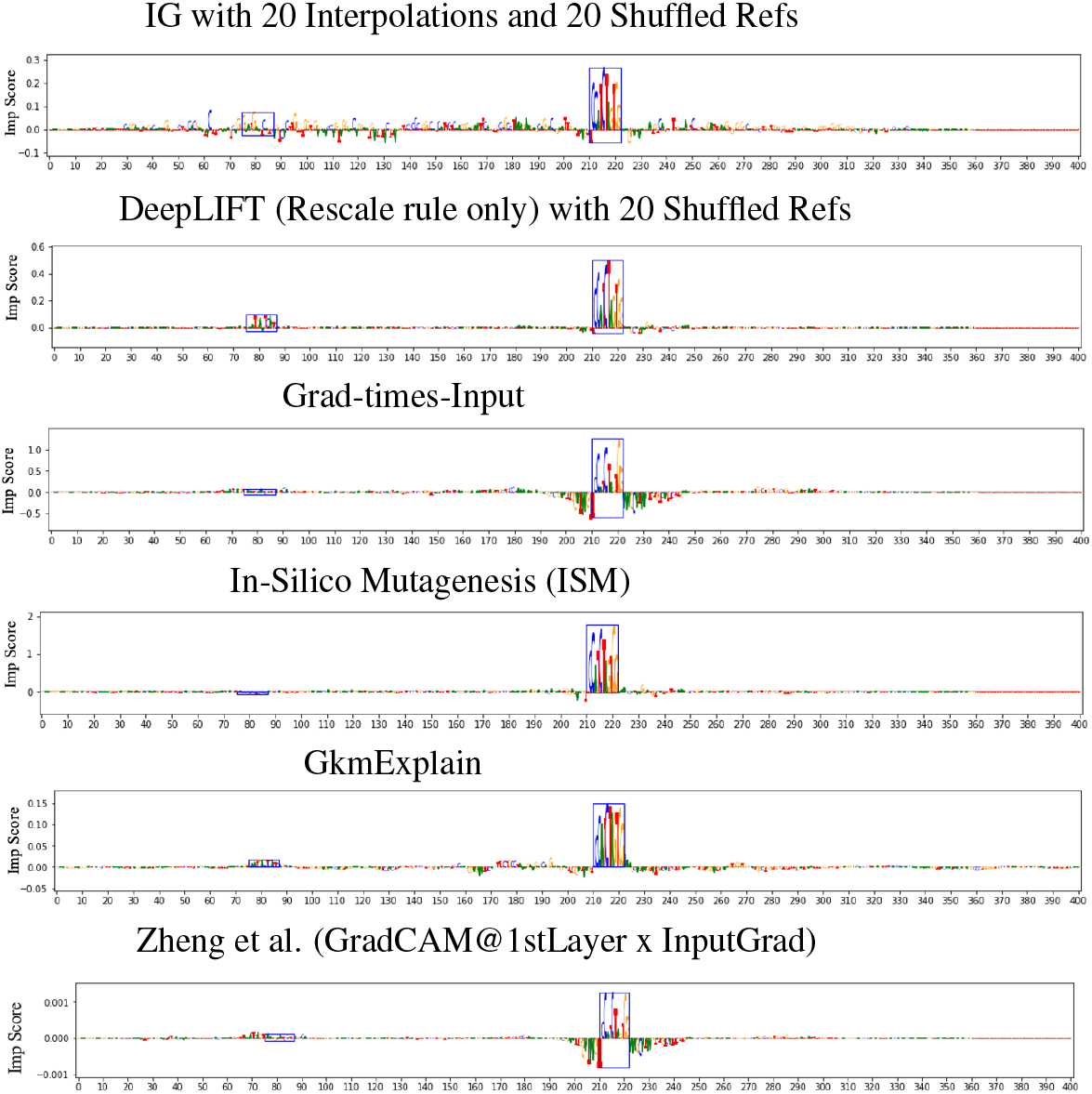
Visualization of importance scores. Shown are scores from GkmExplain and the finetuned Basset model for the simulated sequence corresponding to chr14:64937697-64938097 in A549. Blue boxes represent locations of embedded motifs.

## 4 Discussion

Our results with GradCAM demonstrate the value of realistic simulations; while a GradCAM-based method performed well on the simplified simulation in Zheng et al. [11], it performed relatively poorly on our more complex simulation (particularly for the DeepSEA-based architecture). This is consistent with GradCAM’s use of position-invariant weights 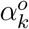. Interestingly, while several refinements of GradCAM exist [30, 31, 32], they all share in common this use of position-invariant weights - for example, LIFT-CAM [31] uses DeepLIFT rather than gradients to compute 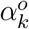. CAM-based methods are popular in computer vision because they tend to be less noisy than computing importance scores directly at the input layer - however, to our knowledge, no work has demonstrated that CAM-based methods outperform simply computing the importance scores at the same layer that the CAM method is applied to. In fact, Rebuffi et al. [27] found gradient-times-activation (which they call “linear approximation”) tends to outperform GradCAM (see Table 3 in their paper), raising questions about whether the benefit of CAM is primarily due to the *layer* at which the importance scores are calculated, as opposed to the postion-invariant channel-weighting approach that CAM uses. Consistent with this hypothesis, we found that DeepLIFT computed at the 1st conv layer sometimes outperforms DeepLIFT computed at the input (**Fig. 4**). This suggests that a promising future direction may be to improve motif identification by combining importance scores computed at multiple different layers, similar to the approaches taken by [11], [27] and [33]. It is also worth investigating *why* importance scores can become worse at highlighting key features when computed closer to the input layer.

We also showed that IG sometimes performs worse than gradient-times-input despite having been developed to address the gradient saturation problem. Our investigations into the output logits suggest the nonlinear interpolation paths used by Jha et al. [17] in the context of splicing codes could prove useful in regulatory genomics as well.

Our experiments found that ISM (conducted at a per-base resolution) performs slightly worse than DeepLIFT at identifying positions within embedded motifs **Fig. 2 & 5**. We suggest two explanations for this: one is that base-resolution ISM may be susceptible to saturation effects [8] for example, mutating a single base may not be enough to substantially disrupt some motif instances; future work could explore the effectiveness of calculating ISM by perturbing sliding windows, rather than one base at a time. Another explanation is that this is a limitation of using a motif discovery method to define ‘ground-truth’ in our simulations; perhaps some motifs identified in the positive set are not in fact bound due to missing contextual features, and ISM correctly reflects this. In this case, a more advanced simulation that explicitly defines contextual features could be a way to go.

Finally, we note that our simulated dataset can be applied not just for bench-marking interpretation, but also for understanding and debugging the learning dynamics of machine learning models applied to regulatory DNA - for instance, are there specific motifs that the models tend to miss? What dataset sizes are needed before a model learns all the motifs that are present? To what extent does pretraining help identify motifs that would have been missed otherwise? Is there an architecture that can learn all the motifs present *without* needing pretraining? We anticipate that this approach may lay the groundwork for future systematic investigations into the training and interpretation of complex machine learning models in regulatory genomics.

## Author Contributions

EP trained the models and ran experiments with guidance from AS. EP implemented the pipelines for generating simulated datasets and computing importance scores, based around code provided by AS. AS conceived of the project and designed experiments. AK provided guidance and feedback. EP, AS and AK wrote the manuscript.

## S1 Supplementary Methods

### S1.1 Enrichment thresholds used for filtering motifs

The set of motifs found by HOMER was further filtered based on the hits identified by FIMO. We set the minimum number of positive motif hits to be 1,000, and also computed an “enrichment score” for the motifs as follows: we assign each motif hit a weight that is inversely proportional to the square of the distance of the hit from the center of the sequence. We then define the “weighted hit density” of a motif in a given set of sequences as the total sum of hit weights over all motif instances in the set divided by the total number of sequences in the set; the “enrichment score” is defined as the weighted hit density on the positive sequences divided by the weighted hit density on the negative sequence. We set the enrichment score threshold to be a minimum of 1.25.

### S1.2 Number of pos/neg sequences in train/test sets

**Figure S6:**
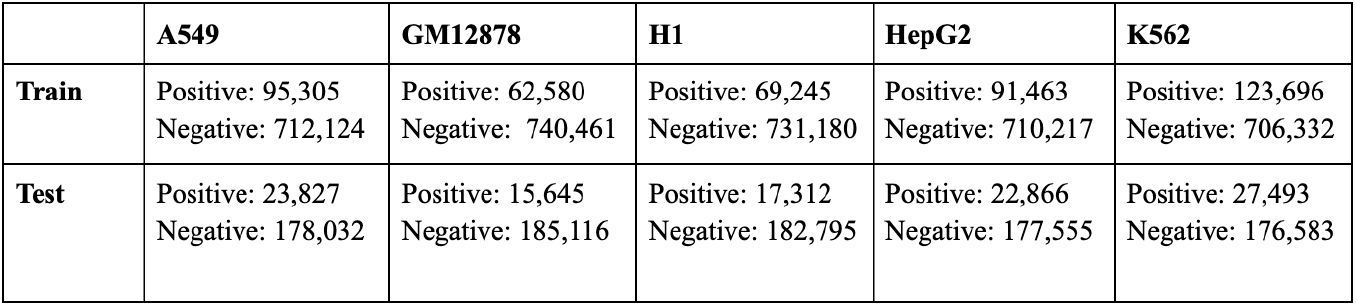
Number of sequences in positive and negative train and test sets in each cell line.

## S2 Supplementary Figures

**Figure S7:**
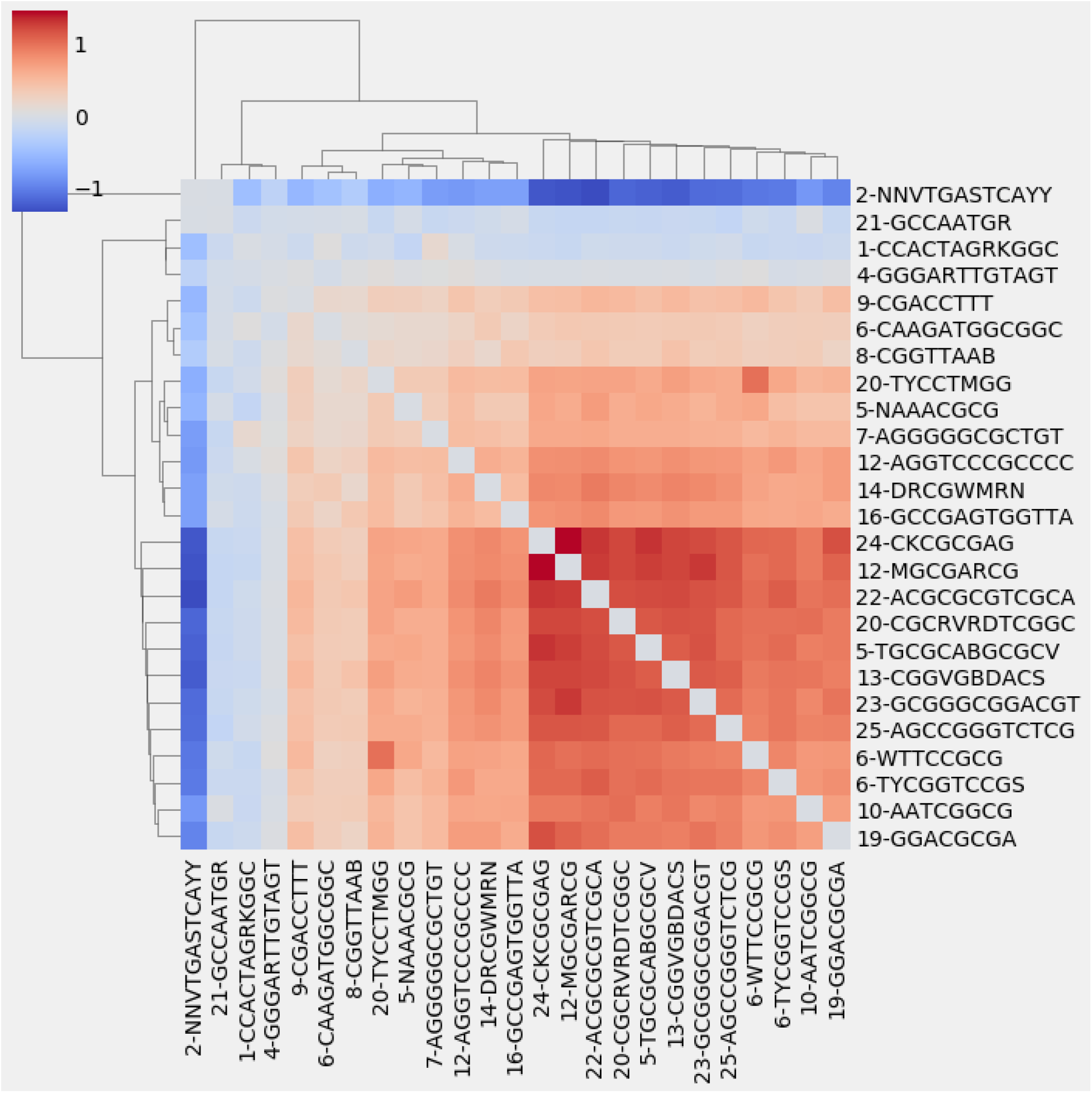
Heatmap of log base 2 fold change (indicated by color intensity) for A549 cell line HOMER accessible sequence motifs called with FIMO. Color indicates fold increase of motif co-occurrence relative to a null where motifs are distributed randomly across sequences.

**Figure S8:**
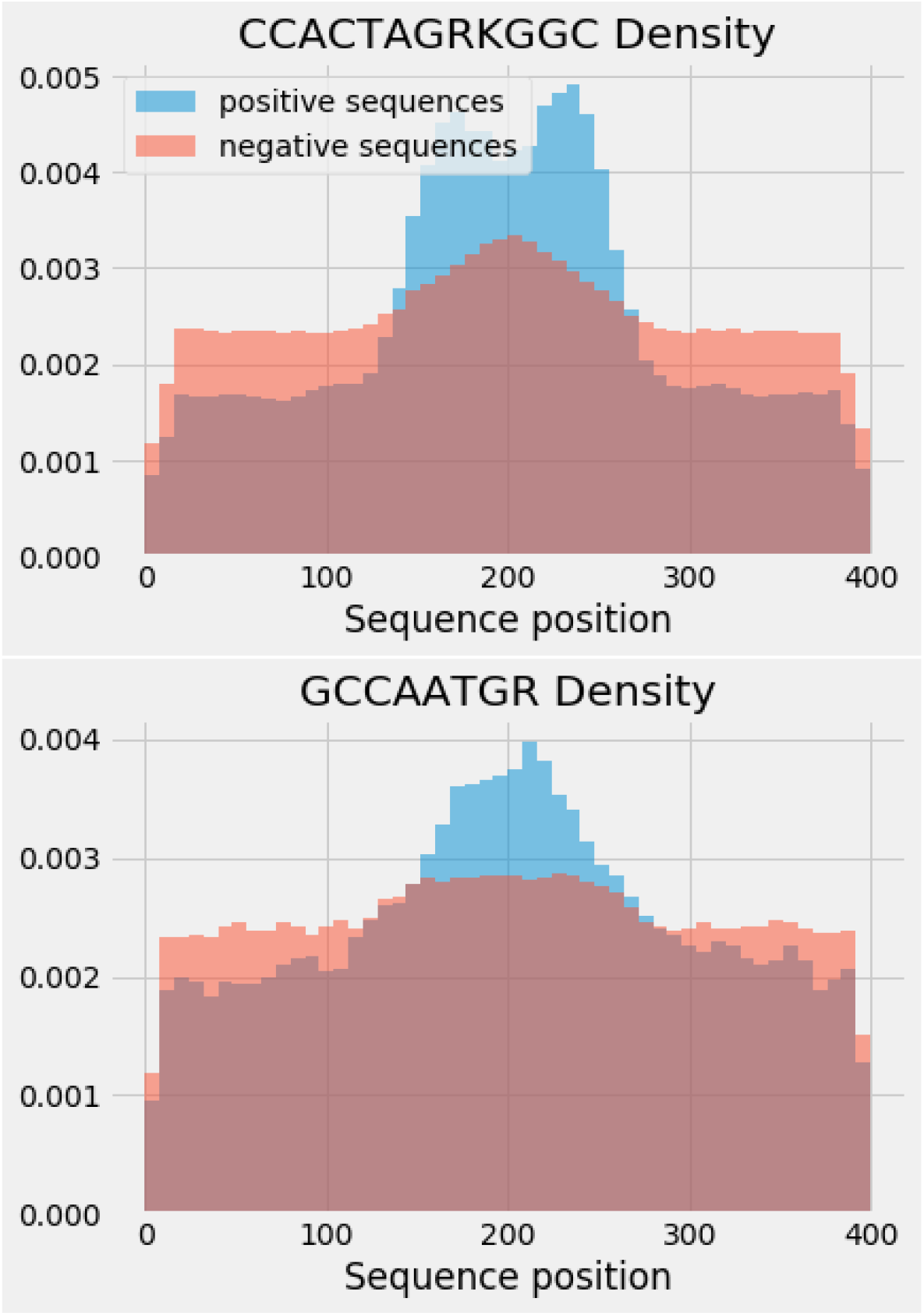
Histograms of positional densities of two A549 motifs.

**Figure S9:**
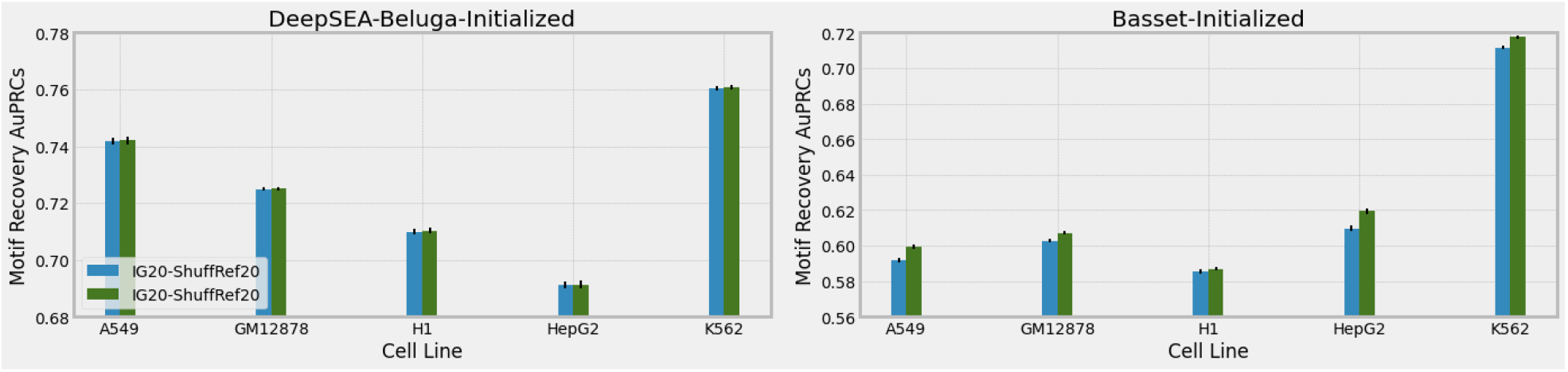
Comparison of number of interpolation points for Integrated Gradients. ‘IG10’ and ‘IG20’ indicated Integrated Gradients with 10 and 20 interpolation points respectively. ‘ShuffRef20’ indicates 20 dinucleotide-shuffled references were used per sequence.

**Figure S10:**
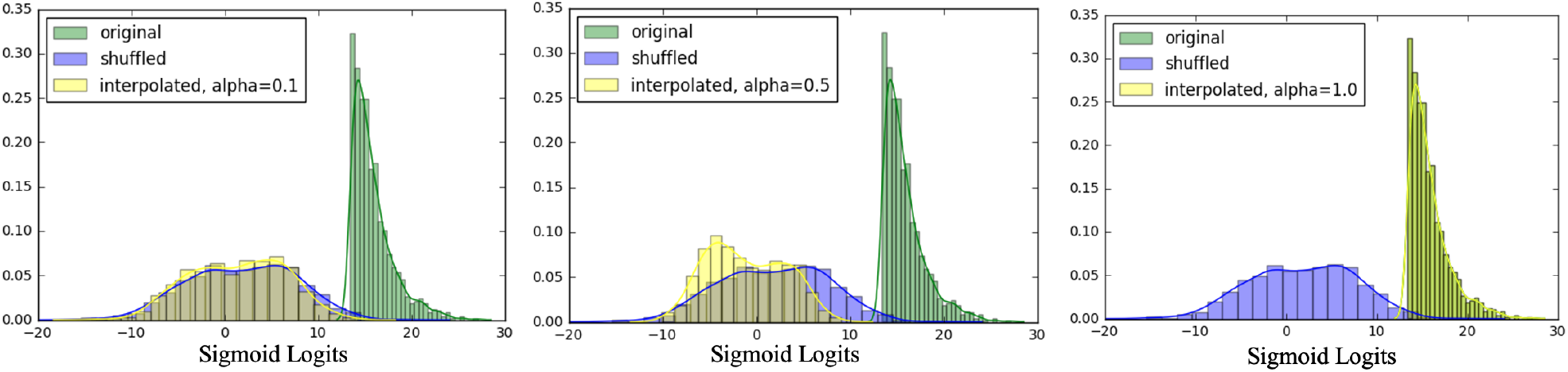
Histograms of sigmoid logits during interpolation between original and dinucleotide shuffled A549 sequences. *α*, which ranges from 0 to 1, indicates the point along the interpolation, with *α* = 0.5 being the halfway point between the shuffled and original sequences.

**Figure S11:**
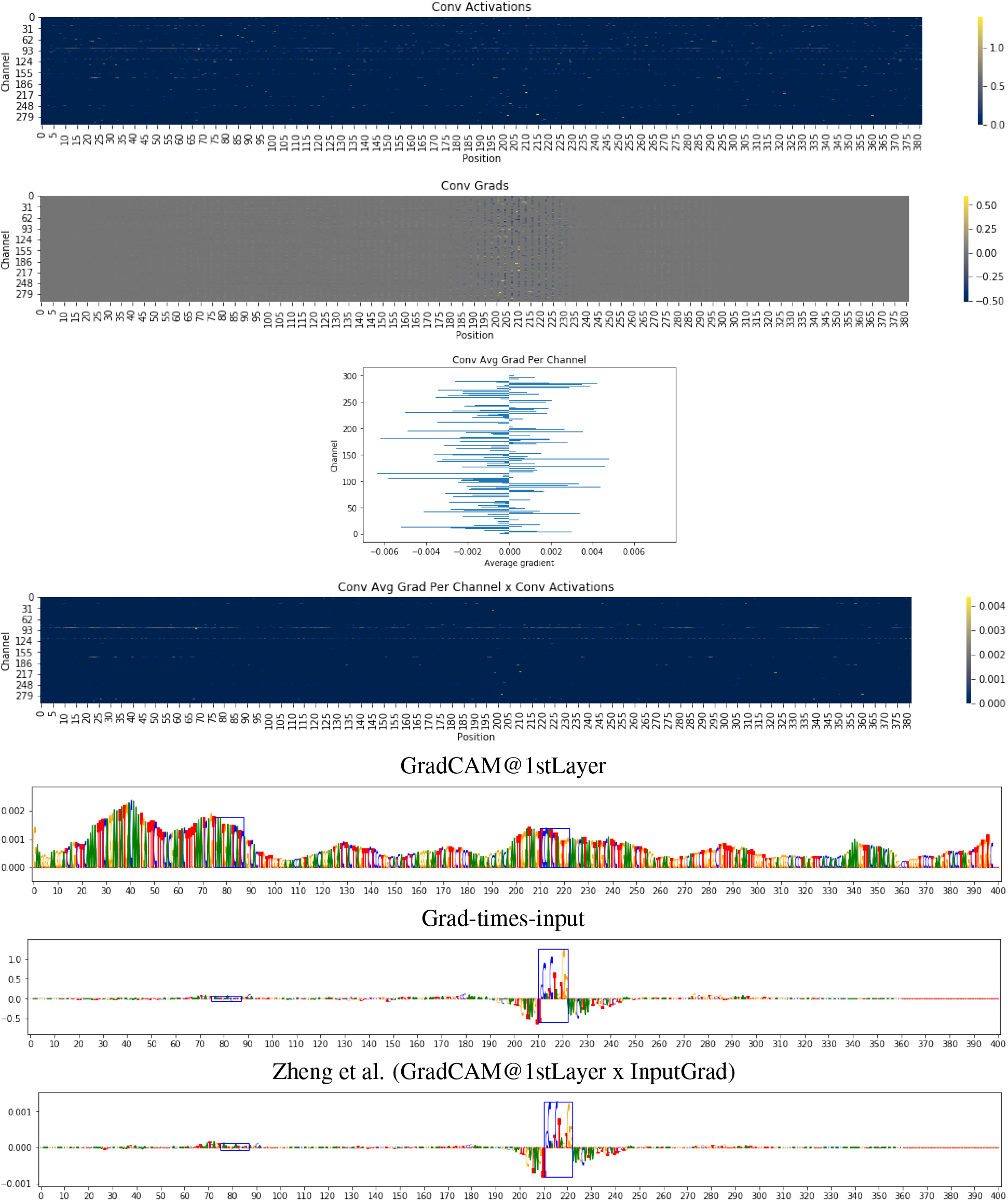
GradCAM discards positional and contextual information (example 1). Each figure depicts a different step in the calculation of importance scores for GradCAM/Zheng et al. Scores are shown for the finetuned Basset model for the simulated sequence corresponding to chr14:64937697-64938097 in A549. Blue boxes represent locations of embedded motifs. Top figure (step 1): heatmap of 1st conv layer activations. Second from top: conv layer gradients. Third from top: gradient averaged across the positional axis for each convolutional channel (there are the *α* values in GradCAM). Fourth from top: product of conv layer activations with the corresponding *α* value for each channel; for ease of visualization, negative values are not shown here. Fifth from the top: GradCAM scores (derived by summing the activation-times-alpha across all channels at each position, taking only positive scores, and projecting the values onto the input layer using the mapping method of Zheng et al.). Sixth from top: gradients on the input bases. Bottom: Zheng et al. scores for the sequence (gradcam scores at the first layer multiplied by the input gradients). Note the loss of positional and contextual information in the fourth step, caused by using the alpha values rather than directly using the conv layer gradients.

**Figure S12:**
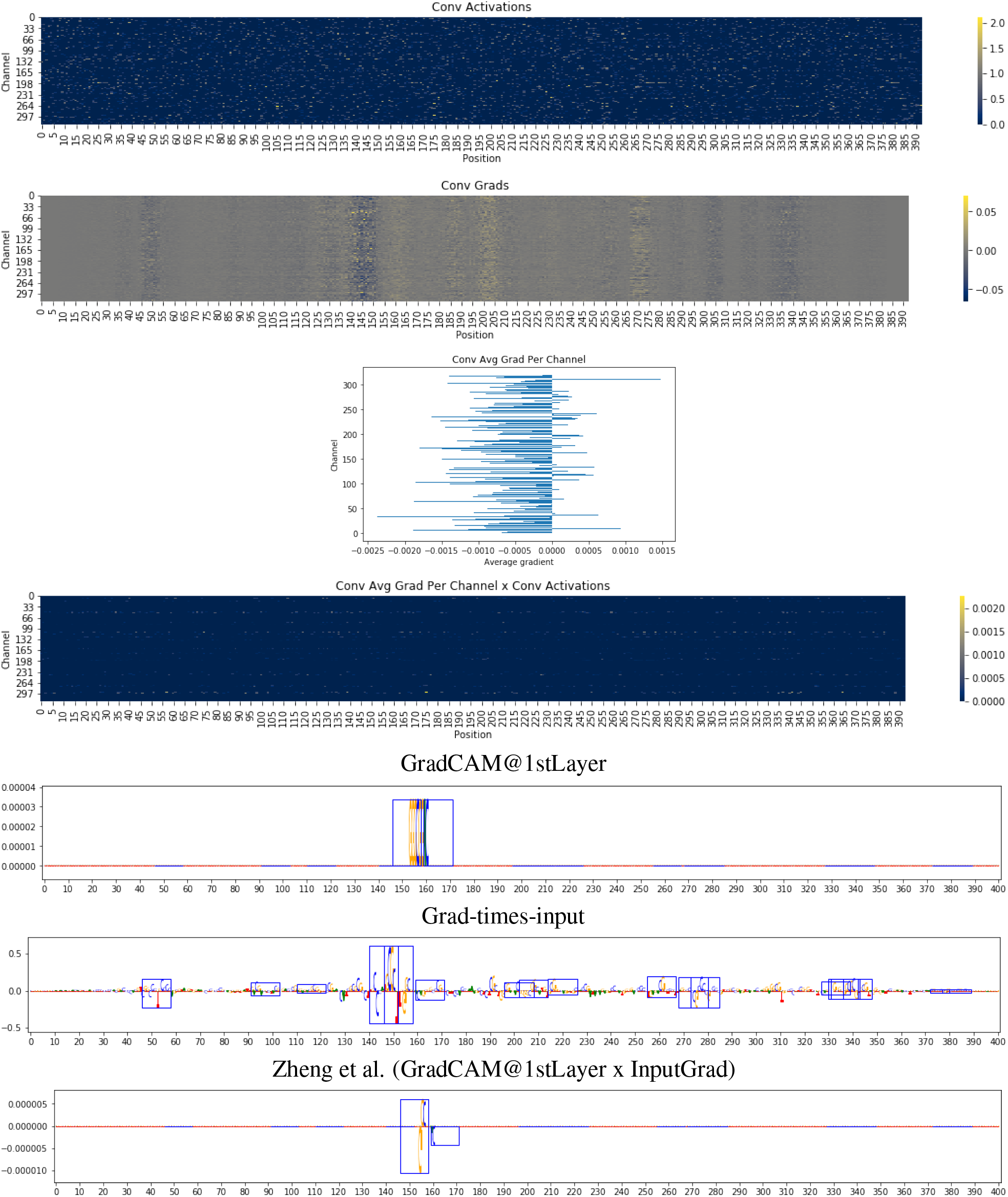
GradCAM discards positional and contextual information (example 2). Similar to **Fig. S11**, but for the finetuned DeepSEA Beluga model for the simulated sequence corresponding to chr19:58543990-58544390 in A549.

**Figure S13:**
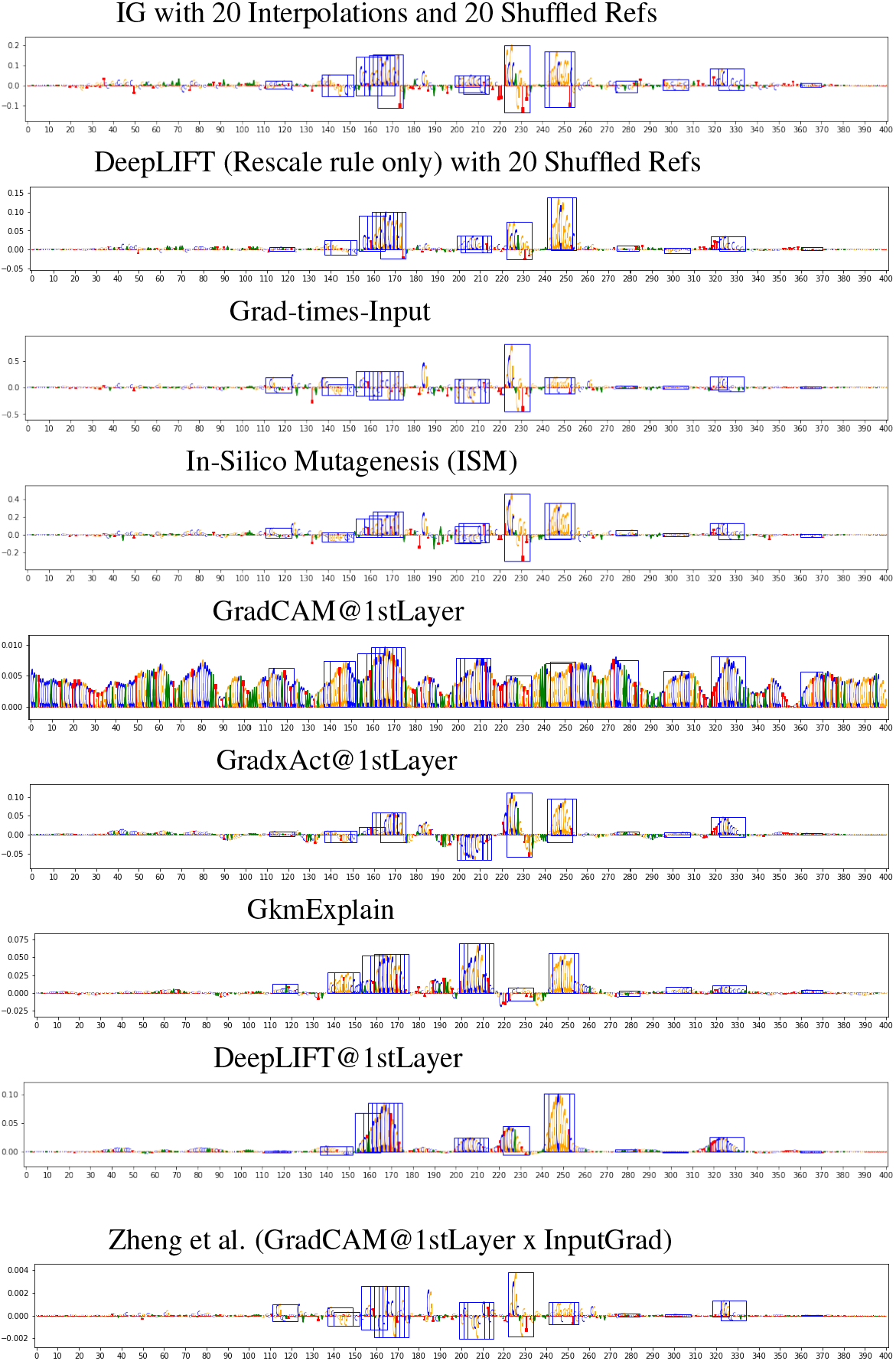
Visualization of importance scores for chr4:74933265-74933665 in A549 for GkmExplain and the finetuned DeepSEA Beluga model. Corresponding scores for the Basset model are in S14. Blue boxes represent locations of embedded motifs. The “GradxAct1stLayer” and “DeepLIFT1stLayer” rows mapped the scores to the input level as described in **Sec. 3.1**. “DeepLIFT1stLayer” used DeepLIFT with the Rescale rule and 20 shuffled references.

**Figure S14:**
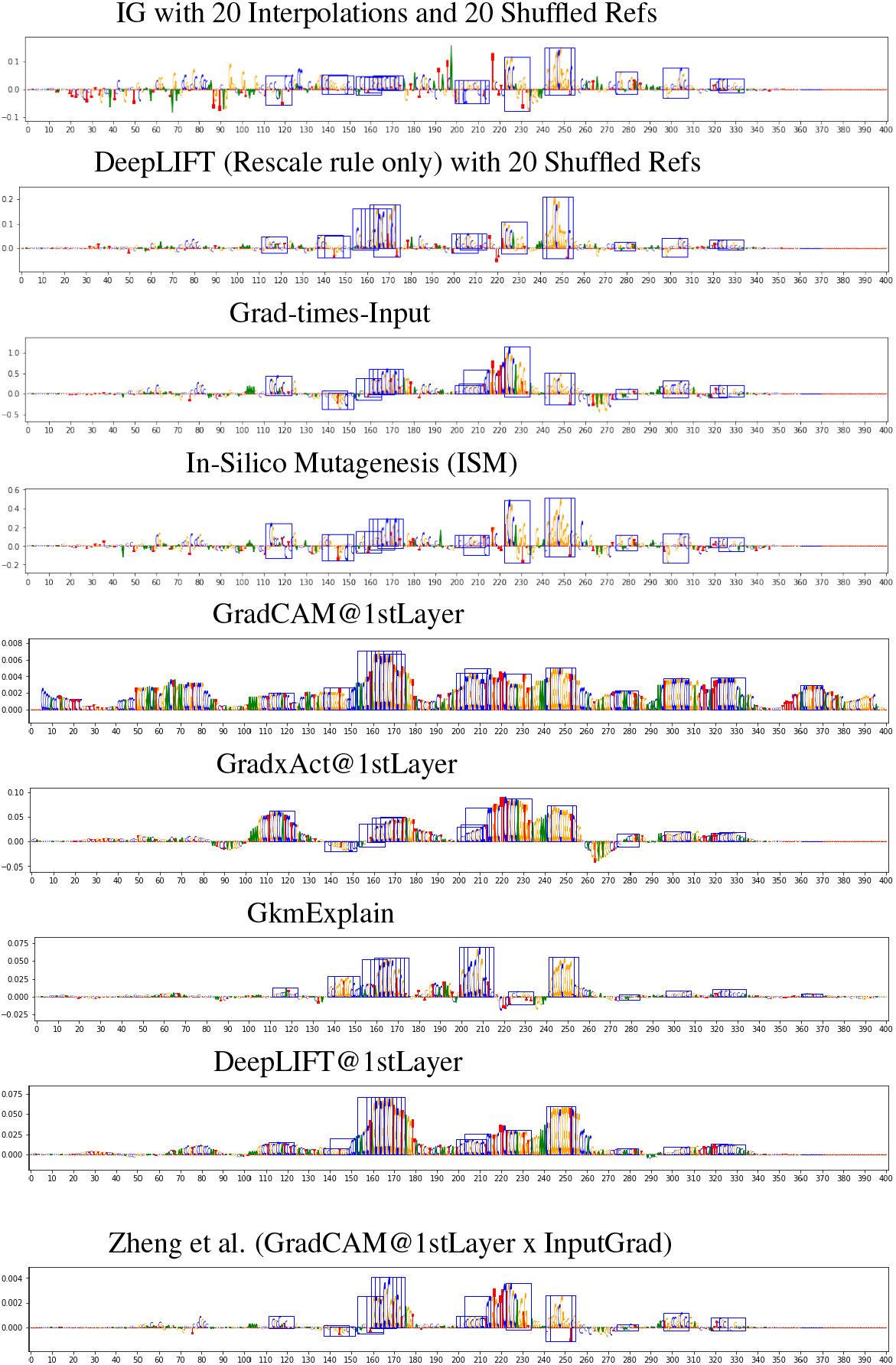
Visualization of importance scores for chr4:74933265-74933665 in A549 for GkmExplain and the finetuned Basset model. Corresponding scores for the DeepSEA Beluga model are in **Fig. S13**. Blue boxes represent locations of embedded motifs. The “GradxAct1stLayer” and “DeepLIFT1stLayer” rows mapped the scores to the input level as described in **Sec. 3.1**. “DeepLIFT1stLayer” used DeepLIFT with the Rescale rule and 20 shuffled references.

## References

[1] Jian Zhou and Olga G Troyanskaya. Predicting effects of noncoding variants with deep learning–based sequence model. Nat. Methods, 12(10):931–934, August 2015.

[2] David R Kelley, Jasper Snoek, and John L Rinn. Basset: learning the regulatory code of the accessible genome with deep convolutional neural networks. Genome Res., 26(7):990–999, July 2016.

[3] Daniel Quang and Xiaohui Xie. FactorNet: A deep learning framework for predicting cell type specific transcription factor binding from nucleotide-resolution sequential data. Methods, 166:40–47, August 2019.

[4] Dongwon Lee. LS-GKM: a new gkm-SVM for large-scale datasets. Bioinformatics, 32(14):2196–2198, July 2016.

[5] Žiga Avsec, Melanie Weilert, Avanti Shrikumar, Sabrina Krueger, Amr Alexandari, Khyati Dalal, Robin Fropf, Charles McAnany, Julien Gagneur, Anshul Kundaje, and Julia Zeitlinger. Base-resolution models of transcription-factor binding reveal soft motif syntax. Nat. Genet., 53(3):354–366, March 2021.

[6] Babak Alipanahi, Andrew Delong, Matthew T Weirauch, and Brendan J Frey. Predicting the sequence specificities of DNA-and RNA-binding proteins by deep learning. Nat. Biotechnol., 33(8):831–838, August 2015.

[7] Ramprasaath R Selvaraju, Michael Cogswell, Abhishek Das, Ramakrishna Vedantam, Devi Parikh, and Dhruv Batra. Grad-cam: Visual explanations from deep networks via gradient-based localization. In Proceedings of the IEEE international conference on computer vision, pages 618–626, 2017.

[8] Avanti Shrikumar, Peyton Greenside, and Anshul Kundaje. Learning important features through propagating activation differences. In Proceedings of the 34th International Conference on Machine Learning, volume 70 of Proceedings of Machine Learning Research, pages 3145–3153. PMLR, 06–11 Aug 2017. URL https://proceedings.mlr.press/v70/shrikumar17a.html.

[9] Mukund Sundararajan, Ankur Taly, and Qiqi Yan. Axiomatic attribution for deep networks. In Proceedings of the 34th International Conference on Machine Learning, volume 70 of Proceedings of Machine Learning Research, pages 3319–3328. PMLR, 06–11 Aug 2017. URL https://proceedings.mlr.press/v70/sundararajan17a.html.

[10] Avanti Shrikumar, Eva Prakash, and Anshul Kundaje. GkmExplain: fast and accurate interpretation of nonlinear gapped k-mer SVMs. Bioinformatics, 35(14):i173–i182, July 2019.

[11] An Zheng, Michael Lamkin, Hanqing Zhao, Cynthia Wu, Hao Su, and Melissa Gymrek. Deep neural networks identify sequence context features predictive of transcription factor binding. Nat Mach Intell, 3(2):172–180, February 2021.

[12] Peter K Koo, Sharon Qian, Gal Kaplun, Verena Volf, and Dimitris Kalimeris. Robust neural networks are more interpretable for genomics. June 2019.

[13] Peter K Koo and Matt Ploenzke. Improving representations of genomic sequence motifs in convolutional networks with exponential activations. Nat Mach Intell, 3(3):258–266, March 2021.

[14] Alexandra Maslova, Ricardo N Ramirez, Ke Ma, Hugo Schmutz, Chendi Wang, Curtis Fox, Bernard Ng, Christophe Benoist, Sara Mostafavi, and Immunological Genome Project. Deep learning of immune cell differentiation. Proc. Natl. Acad. Sci. U. S. A., 117(41):25655–25666, October 2020.

[15] Dongwon Lee, David U Gorkin, Maggie Baker, Benjamin J Strober, Alessandro L Asoni, Andrew S McCallion, and Michael A Beer. A method to predict the impact of regulatory variants from DNA sequence. Nat. Genet., 47(8):955–961, August 2015.

[16] Jian Zhou, Chandra L Theesfeld, Kevin Yao, Kathleen M Chen, Aaron K Wong, and Olga G Troyanskaya. Deep learning sequence-based ab initio prediction of variant effects on expression and disease risk. Nat. Genet., 50(8):1171–1179, August 2018.

[17] Anupama Jha, Joseph K Aicher, Matthew R Gazzara, Deependra Singh, and Yoseph Barash. Enhanced integrated gradients: improving interpretability of deep learning models using splicing codes as a case study. Genome Biol., 21(1):149, June 2020.

[18] Surag Nair, Avanti Shrikumar, and Anshul Kundaje. fastISM: Performant in-silico saturation mutagenesis for convolutional neural networks. October 2020.

[19] Avanti Shrikumar, Peyton Greenside, Anna Shcherbina, and Anshul Kundaje. Not just a black box: Learning important features through propagating activation differences. May 2016.

[20] Scott M Lundberg and Su-In Lee. A unified approach to interpreting model predictions. In I Guyon, U V Luxburg, S Bengio, H Wallach, R Fergus, S Vishwanathan, and R Garnett, editors, Advances in Neural Information Processing Systems 30, pages 4765–4774. Curran Associates, Inc., 2017.

[21] Narine Kokhlikyan, Vivek Miglani, Miguel Martin, Edward Wang, Bilal Alsallakh, Jonathan Reynolds, Alexander Melnikov, Natalia Kliushkina, Carlos Araya, Siqi Yan, and Orion Reblitz-Richardson. Captum: A unified and generic model interpretability library for PyTorch. September 2020.

[22] Marco Ancona, Enea Ceolini, Cengiz Oztireli, and Markus Gross. Towards better understanding of gradient-based attribution methods for deep neural networks. In 6th International Conference on Learning Representations (ICLR 2018), 2018.

[23] Gabriel Erion, Jose Janizek, Pascal Sturmfels, Scott M Lundberg, and Su-In Lee. Improving performance of deep learning models with axiomatic attribution priors and expected gradients. Nature Machine Intelligence, 3(7):620–631, May 2021.

[24] Pascal Sturmfels, Scott Lundberg, and Su-In Lee. Visualizing the impact of feature attribution baselines. Distill, 2020. doi: 10.23915/distill.00022. https://distill.pub/2020/attribution-baselines.

[25] ENCODE Project Consortium. An integrated encyclopedia of DNA elements in the human genome. Nature, 489(7414):57–74, September 2012.

[26] Stephen G Landt, Georgi K Marinov, Anshul Kundaje, Pouya Kheradpour, Florencia Pauli, Serafim Batzoglou, Bradley E Bernstein, Peter Bickel, James B Brown, Philip Cayting, Yiwen Chen, Gilberto DeSalvo, Charles Epstein, Katherine I Fisher-Aylor, Ghia Euskirchen, Mark Gerstein, Jason Gertz, Alexander J Hartemink, Michael M Hoffman, Vishwanath R Iyer, Youngsook L Jung, Subhradip Karmakar, Manolis Kellis, Peter V Kharchenko, Qunhua Li, Tao Liu, X Shirley Liu, Lijia Ma, Aleksandar Milosavljevic, Richard M Myers, Peter J Park, Michael J Pazin, Marc D Perry, Debasish Raha, Timothy E Reddy, Joel Rozowsky, Noam Shoresh, Arend Sidow, Matthew Slattery, John A Stamatoyannopoulos, Michael Y Tolstorukov, Kevin P White, Simon Xi, Peggy J Farnham, Jason D Lieb, Barbara J Wold, and Michael Snyder. ChIP-seq guidelines and practices of the ENCODE and modENCODE consortia. Genome Res., 22(9):1813–1831, September 2012.

[27] Sylvestre-Alvise Rebuffi, Ruth Fong, Xu Ji, and Andrea Vedaldi. There and back again: Revisiting backpropagation saliency methods. In Proceedings of the IEEE/CVF Conference on Computer Vision and Pattern Recognition, pages 8839–8848, 2020.

[28] Asifullah Khan, Anabia Sohail, Umme Zahoora, and Aqsa Saeed Qureshi. A survey of the recent architectures of deep convolutional neural networks. Artificial Intelligence Review, 53(8):5455–5516, December 2020.

[29] Kamil Wnuk, Jeremi Sudol, Kevin B Givechian, Patrick Soon-Shiong, Shahrooz Rabizadeh, Christopher Szeto, and Charles Vaske. Deep learning implicitly handles tissue specific phenomena to predict tumor DNA accessibility and immune activity. iScience, 20:119–136, October 2019.

[30] Aditya Chattopadhyay, Anirban Sarkar, Prantik Howlader, and Vineeth N. Balasubramanian. Gradcam++: Generalized gradient-based visual explanations for deep convolutional networks. CoRR, abs/1710.11063, 2017. URL http://arxiv.org/abs/1710.11063.

[31] Hyungsik Jung and Youngrock Oh. Towards better explanations of class activation mapping. February 2021.

[32] Jeong Ryong Lee, Sewon Kim, Inyong Park, Taejoon Eo, and Dosik Hwang. Relevance-cam: Your model already knows where to look. In Proceedings of the IEEE/CVF Conference on Computer Vision and Pattern Recognition (CVPR), pages 14944–14953, June 2021.

[33] Suraj Srinivas and François Fleuret. Full-gradient representation for neural network visualization. In Advances in Neural Information Processing Systems, volume 32. Curran Associates, Inc., 2019. URL https://proceedings.neurips.cc/paper/2019/file/80537a945c7aaa788ccfcdf1b99b5d8f-Paper.pdf.

